# Trophoblast proliferation, migration, and invasion is regulated by the miR-195/EGFR signaling pathway in recurrent spontaneous abortion patients

**DOI:** 10.1101/630590

**Authors:** Xue Bai, Chunyang Zheng, Na Ren

## Abstract

**OBJECTIVE:** To investigate the effect of the miR-195/EGFR signaling pathway on trophoblasts in patients with recurrent spontaneous abortion, thus providing a clinical basis for the diagnosis and treatment of recurrent spontaneous abortion.

**METHODS:** RT-qPCR, western blot, flow cytometry, CCK8, cell scratch assay, transwell, and a dual Luciferase reporter assay were used to detect changes in the miR-195/EGFR signaling pathways in clinical samples and *in vitro* cultured cells and to explore how these changes affect trophoblasts in affected patients.

**RESULTS:** Expression of miR-195 was elevated in villus tissues of patients with recurrent spontaneous abortion, while the expression levels of EGFR and its downstream genes p38 and AKT phosphorylation were down-regulated. *In vitro* cultured cell experiments showed that miR-195 inhibited the proliferation, migration, and invasion of trophoblast cells. EGFR is a target gene of miR-195, and miR-195 suppresses the expression of EGFR.

**Conclusion:** The miR-195/EGFR signaling pathway regulates the proliferation, migration, and invasion of trophoblast cells, thus playing an important role in recurrent spontaneous abortion.

## 1. INTRODUCTION

Recurrent abortion is defined as a spontaneous abortion that occurs before 20 weeks of gestation three or more times. Recurrent abortion is a common reproductive problem worldwide, affecting up to 1-5% of couples of childbearing age [1–3]. Repeated abortion causes not only great physical damage to pregnant women, but also serious psychological harm. Recurrent abortion is complex and often caused by a combination of factors such as anatomical, genetic, endocrine, infection, immune, and environmental influences [4–6]. More seriously, the cause of recurrent abortion in many patients is unclear [2]. Therefore, deepening the clinical study and experimental theoretical research of recurrent abortion, and determining the pathogenesis of the diagnosis and treatment of recurrent abortion is of great significance.

One of the key events that occurs in pregnancy is the development of a subpopulation of trophoblasts from the trophectoderm implanted in the blastocyst. Trophoblasts have similar proliferation, migration, and invasion capabilities as those of tumor cells, but these capabilities in trophoblasts are regulated by time and space [7]. Trophoblasts are specialized placental cells, and their growth and development are closely related to placental implantation and maternal-fetal interface formation [8].

MicroRNA (miRNA) is a type of non-coding single-stranded RNA molecules of ~23 bp in length, which silences the target gene through RAN interference and participates in post-transcriptional regulation of genes. Dicer is an endoribonuclease that recognizes and cleaves double-stranded RNA. The Dicer enzyme plays an important role in the formation of miRNA and the initial stages of RNA interference. The results of mouse studies demonstrated that females with a conditionally knocked out Dicer gene in the reproductive system had mating behavior but did not produce offspring [9]. Thus, miRNAs are involved in reproductive regulation. Previous studies have also confirmed that multiple miRNAs are involved in reproductive regulation [10]. In the study of eclampsia, miR195 expression changes were related to physiological functions such as trophoblast invasion [11–15]. The epidermal growth factor receptor (EGFR) has tyrosine kinase activity and can be activated by epidermal growth factor (EGF). The EGFR signaling network plays an important role in the growth and maintenance of epithelial tissues. Deregulation or overactivity of the EGFR signaling pathway is common in tumors. In the current study, clinical regulation and *in vitro* culture of cell assays were conducted to investigate the regulation of miR-195/EGFR on trophoblast cells during recurrent spontaneous abortion, thus laying a theoretical foundation for the diagnosis and treatment of recurrent spontaneous abortion.

## 2. Materials and methods

### 2.1 Clinical sample collection

From March 2012 to February 2014, chorionic villus samples from patients with recurrent spontaneous abortion (mean age 30±5 yr; n=20) were collected from the Department of Obstetrics and Gynecology of the General Hospital of Northern Theater Command. We also collected tissue samples from normal pregnant women, (mean age 30±5 years; n=20). Tissue samples were stored at −80 °C. This study was approved by the Ethics Committee of the General Hospital of Northern Theater Command. All patients received written informed consent before the study was enrolled.

### 2.2 Cell culture

The human chorionic trophoblast cell line HTR-8/Svneo was purchased from Shanghai Cellular Library of Chinese Academy of Sciences. RPM-1640 medium (Thermo, Shanghai, China, 11875093) containing 10% fetal bovine serum (Gibco, Australia, 10099141), 100 U/mL penicillin, and 0.1 mg/mL streptomycin (Sigma, Shanghai, China, V900929) was cultured in a 5% CO_2_, 37 °C culture incubator (Thermo Shanghai, China). The medium was changed every other day. When the cell fusion reached 80-90%, it was digested with trypsin (Thermo, Shanghai, China, R001100) and passaged.

### 2.3 Real-time quantitative PCR

Chorionic tissue samples of normal pregnant women and recurrent spontaneous abortion patients were collected clinically. Total RNA from tissue samples was extracted using TRIzol reagent (Thermo, Shanghai, China, 15596026) according to the manufacturer’s instructions. The cDNA was synthesized by reverse transcription using 1 μg of total RNA as a template with a PrimeScript™ RT Reagent Kit with gDNA Eraser (Takara, Beijing, China, RR047A). The SYBR® Premix Ex Taq ^TM^ II kit (Takara, Beijing, China, RR82LR) reagent was used to detect the expression of miR195 by real-time quantitative PCR using the Applied Biosystems 7500 Real-Time PCR System (Thermo, Shanghai, China) with U6 nuclear small RNA as an internal reference. The miR-195 upstream primer was: 5’-ACACTCCAGCTGGGTAGCAGCACAGAAAT-3; the downstream primer was: 5’-TGGTGTCGTGGAGTCG-3. The U6 upstream primer was: 5’-CTCGCTTCGGCAGCACA-3’; the downstream primer was: 5’-AACGCTTCACGAATTTGCGT-3’

### 2.4 Immunoblotting

Total protein was obtained from chorionic tissue of normal pregnant women and recurrent spontaneous abortion patients by cleavage using the RIPA buffer (Sigma, Shanghai, China, V900854). Protein concentration was determined using a BCA protein concentration assay kit (Sigma, Shanghai, China, FP0010). Equal amounts of protein (20 μg) were electrophoresed on a 10% SDS-PAGE gel, electroporated to a PVDF membrane (Sigma, Shanghai, China, IPVH00010), and blocked with 5% BSA in TBST buffer for 1 h at room temperature. After being washed three times with TBST, the corresponding diluted EGFR primary antibody (1:5000, Abcam, Shanghai, China, ab52894), phosphorylated AKT1 primary antibody (1:5000, Abcam, Shanghai, China, ab133458), phosphorylated P38 primary antibody (1:000, Abcam, Shanghai, China, ab178867) and the internal reference β-actin primary antibody (1:5000, Abcam, Shanghai, China, ab8227) were added and incubated overnight at 4 °C with gentle shaking. After washing three times with TBST, the corresponding horseradish peroxidase-labeled secondary antibody (1:5000, CST, Shanghai, China, #7074) was added and incubated for 1 h at room temperature. Color development was performed by adding ECL hypersensitive luminescent solution (Thermo, Shanghai, China, 32132), and gray scale was detected by image laboratory software (Bio-Rad, Hercules, CA, USA) to quantitatively analyze protein expression.

### 2.5 Cell transfection

miR-195 mimic, miR-195 mimic control, miR-195 inhibitor, miR-195 inhibitor control, and luciferase reporter plasmid were synthesized by Gemma Gene Company (Shanghai, China). miR-195 mimic series is 5’-UAGCAGCACAGAAAUGGC-3’; miR-195 mimic control sequence is 5’-UUCUCCGAACGUGUCACGUTT-3’; miR-195 inhibitor sequence is 5’-GCCAAUAUUUCUGUGCUGCUA-3’; miR-195 inhibitor control sequence is 5’-CAGUACUUUUGUGUAGUACAA-3’. Using the primers EGFR 3’ UTR F: 5’ tctagaccactgggcccagaaaggcag 3’ and EGFR 3’ UTR R: 5’ tctagacttaacaatgctgtaggggctctg 3’, the EGFR 3’ UTR fragment was amplified with PCR and inserted into the luciferase reporter vector pGL3 to construct the pGL3-EGFR −3’UTR vector. Transfection was performed using Lipofectamine 2000 (Thermo, Shanghai, China, 11668019).

### 2.6 BrdU labeling and flow detection for cell proliferation

Medium containing 10 μM BrdU (Abcam, Shanghai, China, ab142567) was cultured in the dark for 24 h and BrdU labeling was performed. The medium containing BrdU was removed, washed with PBS, and digested with trypsin, and 1 × 10^6^ cells were collected. Cells were fixed and permeabilized by using Fixation/Permeabilization Solution Kit (BD Biosciences, Shanghai, China, 554714) as instructed. Incubation with 30μg DNase (Sigma, Shanghai, China, D4513) proceeded for 1 h at 37 °C. Cells were washed with Flow Cytometry Staining Buffer (Thermo, Shanghai, China, 00-4222-26) thrice. BrdU-specific antibody (1:100, Abcam, Shanghai, China, ab6326) was added and incubated for 20 min at room temperature. Cells were again thrice washed with Staining Buffer. CY5 conjugated fluorescence secondary antibody (1:500, Abcam, Shanghai, China, ab6565) was added and incubated at room temperature and in the dark for 30 min. Detection was performed using a BD FACSCALIBUR flow cytometer (BD Biosciences, Shanghai, China).

### 2.7 CCK8 detection

The cells were plated at 1 × 10^3^ cells/well in 96-well plates. The CCK8 test kit (Solarbio, Beijing, China, CA1210) was used, and the test was performed at 1, 2, 3, 4, and 5 d.

### 2.8 Cell scratch test

5 × 10^5^ cells and untreated HTR-8/Svneo cells were plated in each well of a 6-well plate. Cell fusion was observed after 12 h. After the cell fusion reached 100%, the head was sterilized and slanted perpendicular to the cell plane and was washed 3 times with PBS to remove the scratched but unattached cells. New medium was added, and culturing continued. The scratch width was observed and recorded at 0 and 12h.

### 2.9 Transwell experiment

Matrigel Matrix (BD Biosciences, Shanghai, China, 356234) diluted 1:5 in serum-free medium was added to the upper chamber of a Transwell chamber (Corning, NY, USA, 3422) with a membrane pore size of 8.0 μm and cultured at 37 °C. At 4 h, the excess fluid was discarded, and the cells were washed with serum-free medium. After digesting HTR-8/Svneo cells with trypsin, a 2 × 105 HTR-8/Svneo cell suspension was prepared using serum-free medium and inoculated into the upper chamber of the Transwell chamber. The lower chamber of the Transwell chamber was filled with complete medium. After 24 h, the Transwell chamber was removed and the Matrigel Matrix at the bottom of the upper chamber and the cells attached thereto were carefully removed with a cotton swab. After washing three times with PBS, the Transwell chamber was placed in 4% paraformaldehyde for 30 min. After 15 min of 0.1% crystal violet staining, the Transwell chamber was placed under a microscope, and 5 scope fields were taken and photographed.

### 2.10 Dual-luciferase reporter experiment

HEK-293T cells were applied into 12-well plates. When the cell fusion reached approximately 90%, miR-195 mimic or miR-195 mimic control was co-transfected with pGL3-EGFR-3’ UTR and internal reference sea cucumber luciferase plasmid phRL-TK, respectively. After 48 h, luciferase activity was measured using a Dual-Luciferase Reporter Assay system (Promega Corporation, Madison, WI, USA)

### 2.11 Data analysis

Data are shown as mean ± standard deviation (SD). Statistical analysis of experimental data was performed using GraphPad prism software. Differences between groups were analyzed by t-test or one-way ANOVA. P < 0.05 was considered statistically significant.

## 3. RESULTS

### 3.1 Up-regulation of miR-195 in villus tissue in patients with recurrent spontaneous abortion

RT-qPCR indicated that the expression of miR-195 in the villus trophoblast cells of patients with recurrent spontaneous abortion was significantly higher than that of normal pregnant women (Fig. 1).

**Figure 1.**
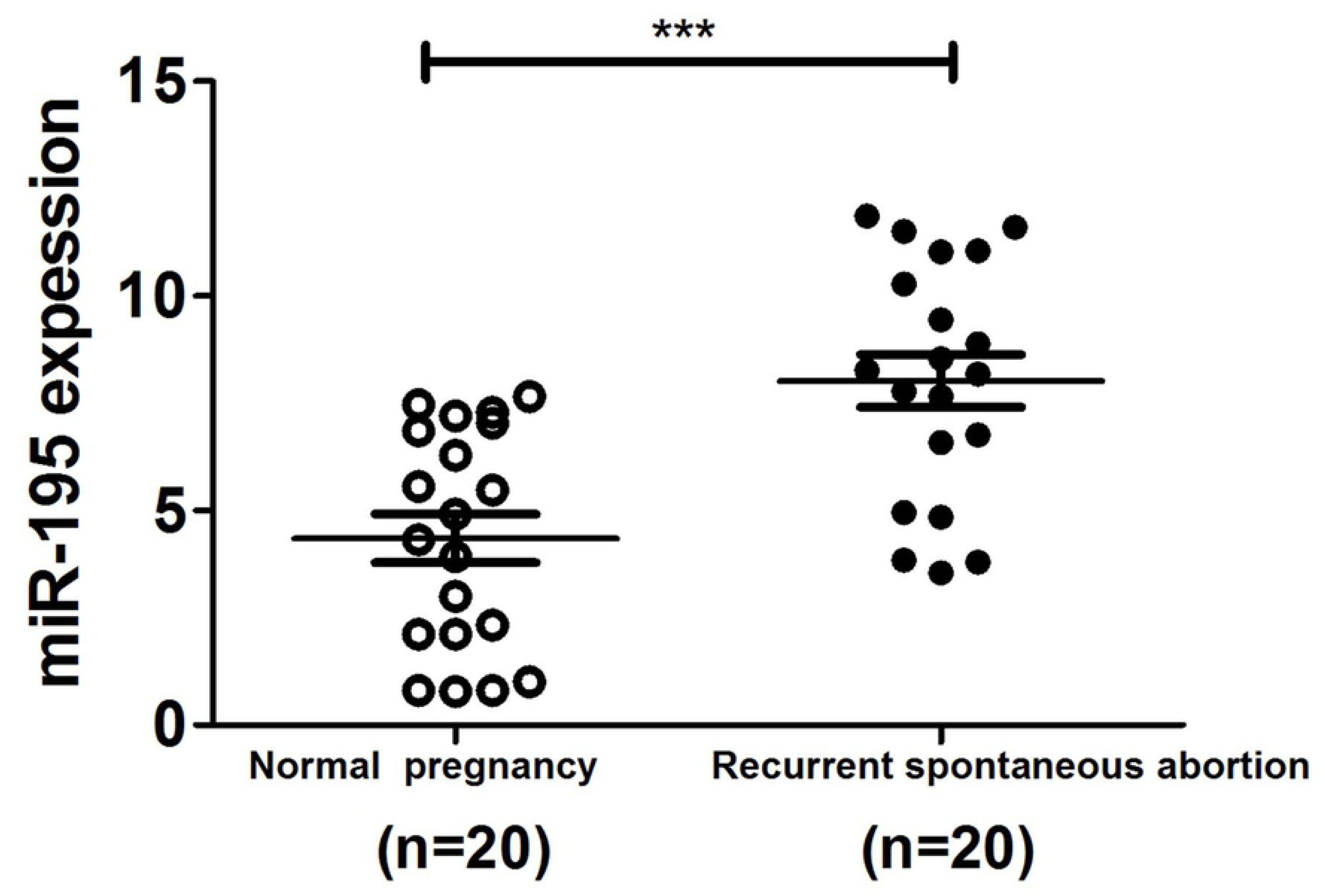
Up-regulation of miR-195 expression in villus tissues of patients with recurrent spontaneous abortion. RT-qPCR determined that the expression of miR-195 in villus tissues of patients with recurrent spontaneous abortion was higher than that in normal pregnant women.

### 3.2 EGFR expression and the downstream signaling pathway are inhibited in patients with recurrent spontaneous abortion

Western blot indicated that the expression of EGFR in villus tissues of patients with recurrent spontaneous abortion was significantly lower than that of normal pregnant women. Moreover, the phosphorylation levels of EGFR-associated PI3K signaling pathway and MAPK signaling pathway markers AKT and p38 in villus tissues of these patients were also significantly lower.

**Figure 2.**
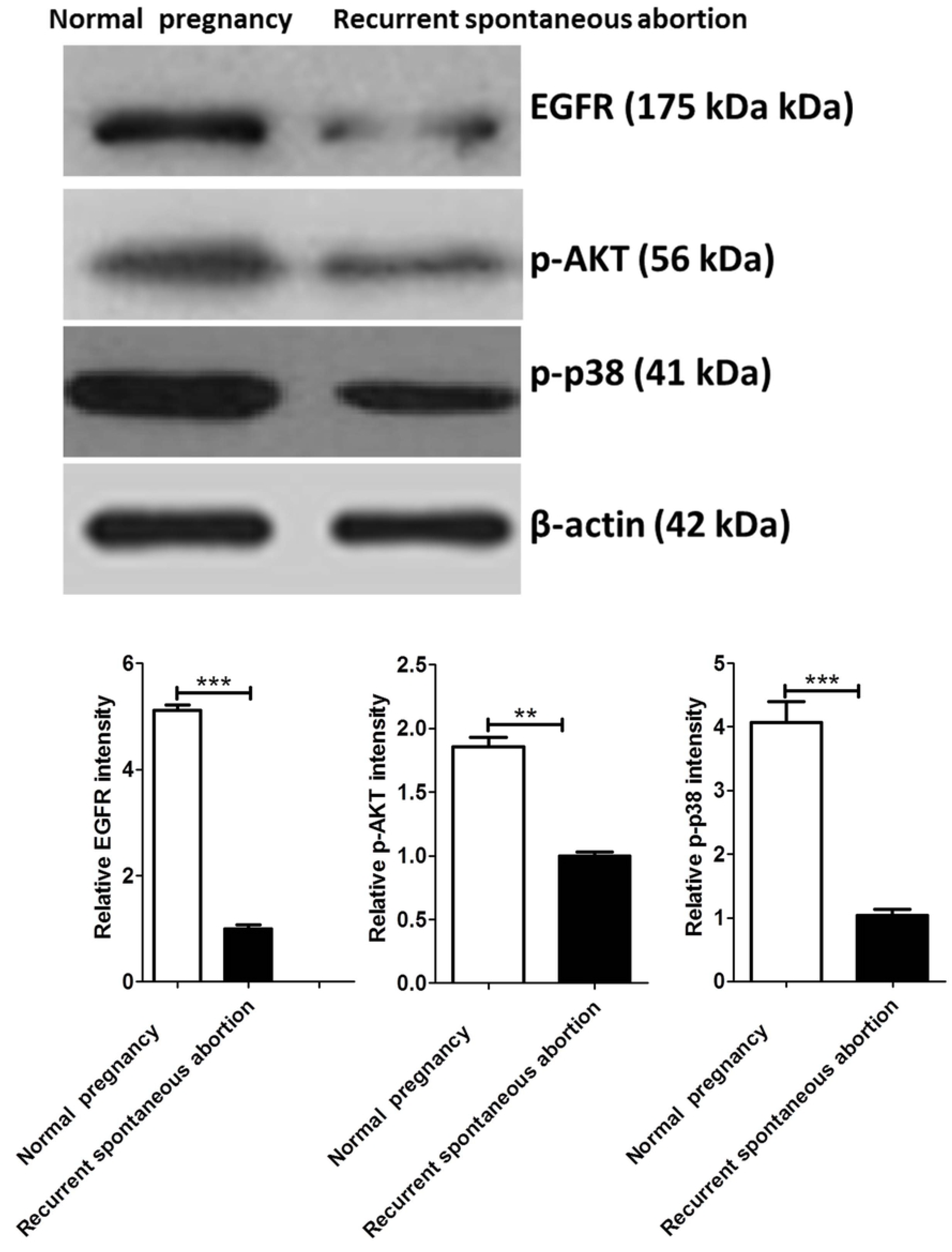
Expression of EGFR in villus tissues and the inhibited EGFR downstream signaling pathway activation in patients with recurrent spontaneous abortion. Western blot analysis showed that the expression of EGFR in villus tissues and the levels of AKT and p38 phosphorylation downstream of EGFR in patients with recurrent spontaneous abortion were significantly lower than those in normal pregnant women. Each test was repeated three times independently and the data is expressed as mean ± SD, *p < 0.05, **p < 0.01, ***p < 0.001.

### 3.3 miR-195 restricts the proliferation, migration, and invasion of trophoblasts

After the miR-195 mimics were transfected with HTR-8/Svneo cells, the effects of miR-195 on the proliferation, migration, and invasion of trophoblast cells were detected. Cell proliferation detected by flow cytometry and CCK8 test results showed that miR-195 inhibited the proliferation of trophoblast cells. The scratch test results suggested that miR-195 hinders the migration of trophoblast cells. Transwell results revealed that miR-195 reduces the invasion of trophoblast cells.

**Figure 3.**
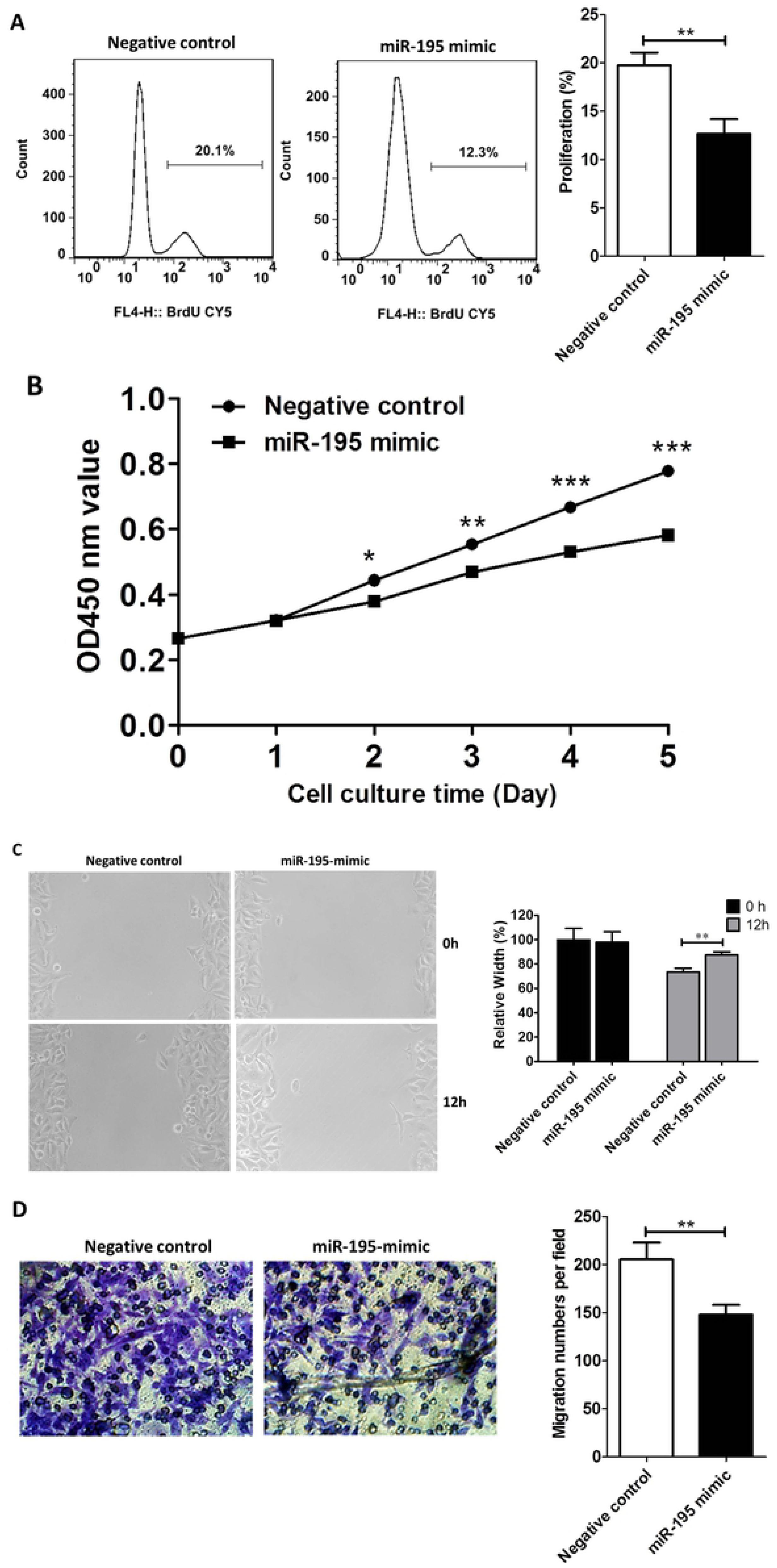
miR-195 inhibits the proliferation, migration, and invasion of trophoblasts. (A) Flow cytometry analysis demonstrated reduced trophoblast proliferation with the miR-195 mimic. (B) CCK8 assay revealed inhibited trophoblast proliferation with the miR-195 mimic. (C) Cell scratch test results indicated reduced proliferation and migration with the miR-195 mimic. (D) Transwell results showed inhibited trophoblast invasion with the addition of the miR-195 mimic,

### 3.4 miR-195 inhibits trophoblast proliferation and migration by regulating EGFR expression

To verify whether miR-195 directly regulates EGFR, we constructed the pGL3-EGFR-3’UTR vector by inserting the EGFR 3’UTR region into the luciferase reporter vector pGL3, termed pGL3-EGFR-3’UTR, and then co-transfected with the miR-195 mimic to perform the dual-luciferase activity assay. The results indicated that miR-195 mimic significantly suppressed the luciferase activity of pGL3-EGFR-3’UTR.

**Figure 4.**
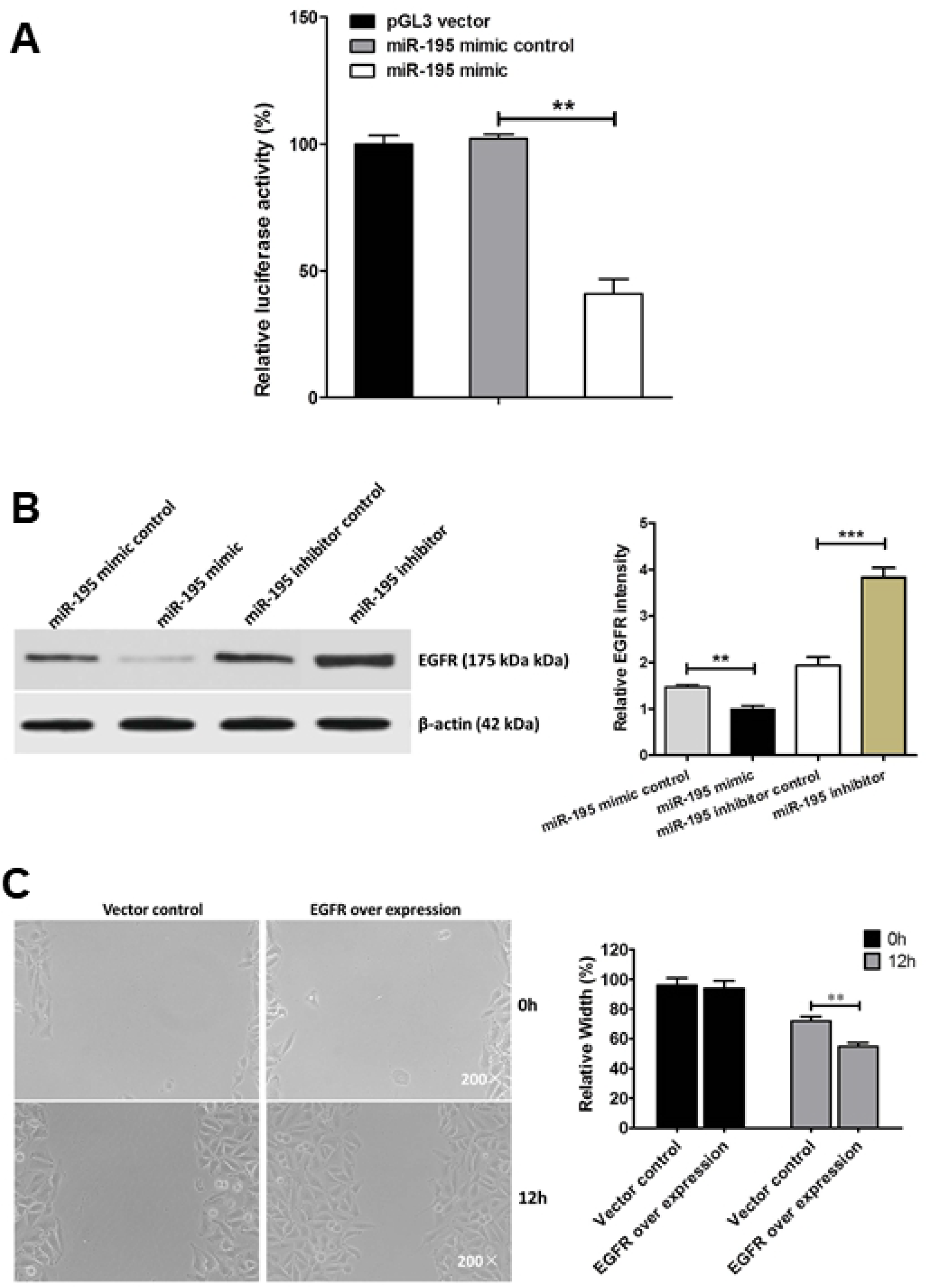
miR-195 inhibits trophoblast proliferation and migration by regulating EGFR expression. (A) A dual-luciferase reporter gene assay indicated that the miR-195 mimic significantly inhibited the luciferase activity of pGL3-EGFR-3’UTR. (B) Western blot analysis showed that the addition of the miR-195 mimic inhibited the expression of EGFR in trophoblast cells, but the addition of miR-195 inhibitor reversed the effect of miR-195 and promoted the expression of EGFR in trophoblast cells. (C) Cell scratch test results revealed that overexpression of EGFR promotes trophoblast proliferation and migration. Each test was repeated three times independently and the data was expressed as mean ± SD, *p < 0.05, **p < 0.01, ***p < 0.001.

To further demonstrate that miR-195 regulates trophoblastic EGFR protein expression, we transfected miR-195 mimic, miR-195 inhibitor, and their corresponding control vectors into HTR-8/Svneo cells. Western blot analysis showed that the miR-195 mimic inhibited EGFR expression, while the miR-195 inhibitor control promoted EGFR expression.

To confirm that EGFR is involved in the regulation of proliferation and invasion of trophoblasts, we transfected EGFR overexpression vectors and its control vectors into HTR-8/Svneo cells and performed scratch assays. The proliferation and invasion abilities of trophoblast cells were significantly increased after EGFR overexpression compared with those of the control.

## 4. DISCUSSION

Recurrent spontaneous abortion seriously jeopardizes the physical and mental health of pregnant women. However, its etiology is complicated, and is poorly understood in many patients because it may be caused by a single factor or a combination of multiple potential factors. Changes in miR-195 expression are often associated with disease progression. Studies have shown that miR-195 expression is down-regulated in a variety of solid tumors such as gastric and colorectal cancers [16–18]. Conversely, some studies have shown that up-regulation of miR-195 expression can discourage tumor cell migration and invasion [19]. The ability of trophoblasts to proliferate, migrate, and invade also plays an important role in the development of the placenta. In studies of eclampsia, miR-195 expression changes were demonstrated to affect embryonic trophoblast infiltration. Our results show that miR-195 expression is increased in villus trophoblast cells in patients with recurrent spontaneous abortion compared to its expression in normal pregnant women. *In vitro* cultured cell experiments determined that miR-195 inhibited the proliferation, migration, and invasion of trophoblast cells. Moreover, here we found that EGFR expression and downstream phosphorylation of p38 and AKT in the villus trophoblast cells of recurrent spontaneous abortion patients were significantly decreased. Both p38 and AKT are important signaling pathway molecules downstream of EGFR. Previous studies have shown that the EGFR signaling pathway plays an important role in tumorigenesis because it regulates tumor cell proliferation, migration, and inhibition [20–22].

The results of hepatocellular carcinoma studies show that EGFR is possibly a target gene of miR-195 and that miR-195/EGFR has a role in regulating cancer cell proliferation, migration, and invasion [23]. Therefore, we speculate that the same regulation is also present in trophoblasts. To confirm this inference, we used a miR-195 inhibitor and performed a dual-luciferase reporter gene assay and found that miR-195 can effectively down-regulate the expression of EGFR in trophoblast cells cultured *in vitro*. After the inhibition of miR-195, the expression of EGFR in cultured trophoblast cells was significantly increased. The dual-luciferase reporter gene results showed that miR-195 acts directly on the 3’ UTR region of EGFR. Cell scratch test results demonstrated that overexpression of EGFR can limit the migration of trophoblast cells. In summary, EGFR is a target gene of miR-195, and miR-195 inhibits the expression of EGFR. The miR-195/EGFR signaling pathway regulates the proliferation, migration, and invasion of trophoblast cells and thus plays an important role in the recurrent flow.

